# Norovirus-mediated modification of the translational landscape via virus and host-induced cleavage of translation initiation factors

**DOI:** 10.1101/060772

**Authors:** Edward Emmott, Frederic Sorgeloos, Sarah L. Caddy, Surender Vashist, Stanislav Sosnovtsev, Richard Lloyd, Kate Heesom, Ian Goodfellow

## Abstract

Noroviruses produce viral RNAs lacking a 5’ cap structure and instead use a virus-encoded VPg protein covalently linked to viral RNA to interact with translation initiation factors and drive viral protein synthesis. Norovirus infection results in the induction of the innate response leading to interferon stimulated gene (ISG) transcription. However the translation of the induced ISG mRNAs is suppressed. Using a novel mass spectrometry approach we demonstrate that diminished host mRNA translation correlates with changes to the composition of the eukaryotic initiation factor complex. The suppression of host ISG translation correlates with the activity of the viral protease (NS6) and the activation of cellular caspases leading to the establishment of an apoptotic environment. These results indicate that noroviruses exploit the differences between viral VPg-dependent and cellular cap-dependent translation in order to diminish the host response to infection.

## Introduction

Noroviruses are the causative agent of the majority of human viral gastroenteritis cases in the developed world (Lopman, 2015). Globally, they are responsible for an estimated 200,000 deaths in children under the age of five in developing countries, and in developed countries noroviruses are a major burden on national healthcare infrastructure due to closed wards and economic costs (Lopman, 2015). Noroviruses are small, single-stranded, positive-sense RNA viruses best known for infecting humans, but several animal-specific noroviruses have also been identified (Karst et al., 2014; Thorne and Goodfellow, 2014).

As members of the *Calidviridae*, noroviruses use a virus-encoded VPg protein in place of a 5’ cap structure to recruit eukaryotic initiation factors and direct translation of viral RNAs (Chaudhry et al., 2006a; Chung et al., 2014; Goodfellow et al., 2005; Leen et al., 2016). The norovirus non-structural proteins are generated by the cleavage of a large polyprotein by the viral protease NS6 (Fig. 1A) (Belliot et al., 2003; Sosnovtsev et al., 2006) whereas the structural proteins VP1 and VP2 are produced from a subgenomic mRNA produced during replication (Yunus et al., 2015). In the case of murine norovirus (MNV), the only norovirus that can undergo efficient replication in cell culture, a single overlapping reading frame encoding a virulence factor is also present within the VP1 coding region (McFadden et al., 2011). Members of the *Norovirus* genus appear distinct from other caliciviruses in that VPg interacts directly with the scaffolding protein eIF4G (Chung et al., 2014; Leen et al., 2016), with this representing the key interaction for viral translation, rather than the cap-binding protein eIF4E (Chaudhry et al., 2006b; Goodfellow et al., 2005; Hosmillo et al., 2014), a further departure from the usual cap-dependent mechanism of protein translation. In addition we have also shown that norovirus infection causes eIF4E phosphorylation, which may lead to the preferential translation of distinct subsets of cellular mRNAs (Royall et al., 2015). Other viruses utilize discrepancies between cellular and viral translation to either enable more efficient translation of viral mRNA in the presence of vastly more abundant cellular mRNA (Firth and Brierley, 2012), or to inhibit the translation of cellular mRNA inhibitory to viral infection (Mohr and Sonenberg, 2012).

**Figure 1.**
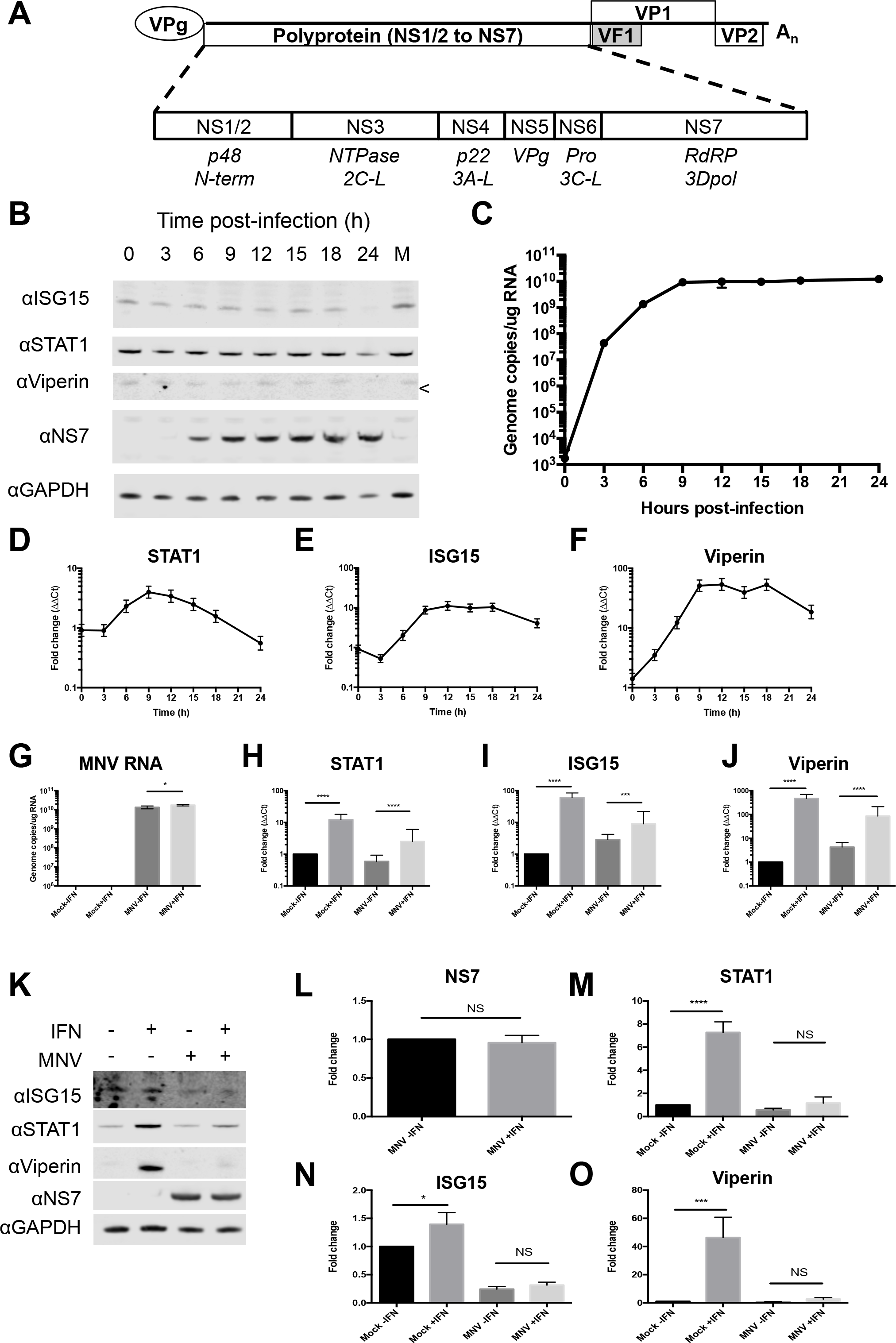
A defect in ISG protein synthesis, but not mRNA induction is observed during norovirus infection. A) The four previously described murine norovirus open reading frames are shown (McFadden et al., 2011). The NS1-7 nomenclature for the mature peptides generated from ORF1 (described in (Sosnovtsev et al., 2006)) is used throughout. B) Western blot and C) RNA levels for norovirus proteins and the ISGs STAT1, ISG15 and viperin (D-F) for a high multiplicity of infection timecourse are shown. On the viperin western blot, the correct position for viperin is indicated with ‘<’. Next, samples were harvested at 18h post-infection following treatment with interferon or mock culture supernatant. qRT-PCR of norovirus and ISG mRNA levels following interferon treatment (G-J) and representative western blots are shown in K), with densitometry analysis of protein levels in L-O) Error bars show standard deviation. Statistical analysis was performed by one-way ANOVA, with the exception of figure L) which was performed by t-test. (*=<0.05, **=<0.01, ***=<0.001, ****=<0.0001).

Studies using MNV, the most common model for the study of norovirus-host interactions, have also confirmed the essential role that the innate response plays in the regulation of norovirus infection and pathogenesis. MNV was discovered in RAG/STAT1^-/-^ mice (Karst et al., 2003), and more recent studies have shown that norovirus infection can be controlled and cleared through interferon λ, with interferon α and β protecting the host from systemic infection even in the absence of adaptive immunity (Mumphrey et al., 2007; Nice et al., 2015). Noroviruses are thought to use several mechanisms to combat the immune response to infection (Roth and Karst, 2015). These include disrupting protein export by the human norovirus NS1/2 (Ettayebi and Hardy, 2003; Fernandez-Vega et al., 2004) or NS4 proteins (Sharp et al., 2012, 2010) as well as diminishing interferon-stimulated gene (ISG) induction through the activity of the MNV virulence factor VF1 protein (McFadden et al., 2011). Despite these mechanisms, the interferon response is activated during both natural infection in humans (Newman et al., 2015) or in mice infected with MNV (Mumphrey et al., 2007) leading to the induction of ISG transcription. Notwithstanding this, we and others have observed that ISG gene induction often does not correlate with the resulting levels of the induced protein (McFadden et al., 2011; Waugh et al., 2014), suggesting a posttranscriptional regulatory mechanism is also involved in the control of the innate response.

We investigated the posttranscriptional regulation of ISG mRNA translation during norovirus infection. Quantitative proteomics was used to identify specific changes to levels or activity of translation initiation factors within norovirus-infected cells. We observed that alterations to the translatome due to norovirus infection were caused by both direct viral and cellular response mechanisms, resulting in the specific reduction in translation of cellular mRNAs. Inhibition of these modifications lead to the restoration of ISG translation and an impact on viral replication, indicating that norovirus infection limits the ability of the innate immune response to combat infection by posttranscriptional regulation of induced mRNAs.

## Results

### Induction of the innate immune response is a late event in norovirus infection

To examine the kinetics of the induction of the innate response during norovirus infection, we examined the levels of ISG mRNA and proteins produced during highly synchronized infection of immortalized macrophage cells (Fig. 1B,C). The levels of representative ISG mRNAs, STAT1, ISG15 and viperin, peaked at around 9 hours post infection and remained at high levels in infected cells (Fig 1D-F). In contrast, levels of the corresponding ISG proteins remained largely unchanged (Fig. 1B).

To examine if norovirus replication affected the ability of cells to respond to interferon we examined the effect of interferon treatment during ongoing norovirus replication (Fig. 1G). Treating infected cells with interferon 6h postinfection lead to robust levels of ISG mRNA induction whilst having a minimal effect on MNV replication (Fig. 1G–J). The levels of ISG mRNA produced following IFN treatment of infected cells were slightly, but significantly (p≤0.001), decreased when compared to uninfected cells (Fig. 1H–J). However, despite robust levels of ISG mRNA transcription, a clear defect in ISG protein production was observed in norovirus infected cells following IFN treatment (Fig. 1K–O).

### Norovirus infection leads to a translational bias

To examine the impact of norovirus infection on global cellular translation, *de.novo* protein synthesis was monitored during synchronized infection by ^35^S methionine pulse-labeling. A clear shift in the translation profile was observed in both RAW264.7 (Fig. 2A) and BV-2 (Fig. 2B) cell lines from 9h post infection, though this fell short of the host shutoff observed in picornavirus or feline calicivirus infection (Gradi et al., 1998; Kuyumcu-Martinez M. et al., 2004). In line with previous polysome analysis performed on norovirus infected cells (Royall et al., 2015), a modest loss of polysomes was observed in RAW264.7 cells with a corresponding increase in the 80s monosome peak (Fig. 2C). A much more apparent reduction in polysome formation occurred during norovirus infection of BV-2 cells (Fig. 2D). Radio-immunoprecipitation of pulse-labeled proteins from norovirus-infected cells confirmed that viral translation was ongoing whilst cellular translation is hindered at 12 hours post infection (Fig. S1A). MNV infection of BV-2 cells follows a similar course to that observed in RAW 264.7 cells (Fig. S1B-D). The loss of polysomes, and increase in monosomes, is characteristic of a defect in translation initiation (Strnadova et al., 2015). Under the conditions used 40s and 60s subunits often associate forming RNA-free 80s monomers, which can be dissociated into free subunits under high salt conditions, whilst RNA-associated ribosomes remain intact. When polysomes were fractionated under high salt conditions the 80s peak dissociated into 40s and 60s monomers confirming that the 80s monomers were not RNA-associated (Fig. S2A,B). A frequently observed mechanism of regulating cellular translation inhibition during viral infection involves the phosphorylation of eIF2α (Clemens, 2001). While modest levels of eIF2α phosphorylation were observed during MNV infection (Fig. S2C), the kinetics of phosphorylation varied in a cell type specific manner and had a poor temporal association with the observed effect on the translation profile, particularly in BV-2 cells. Based on these observations, and that eIF2α phosphorylation would also be anticipated to be inhibitory for norovirus translation, which was not in evidence, we concluded that under the experimental conditions used here, eIF2α phosphorylation did not significantly contribute to the translational bias observed during infection. Studies on pox virus infected cells have suggested that the sequestration of sites of active translation to centers of virus replication causes a translational bias (David et al., 2012). This possibility was investigated using puromycylation to visualize active sites of protein synthesis by covalent linkage of puromycin to newly synthesized peptides as described (David et al., 2012). Puromycylation of mock or infected BV-2 cells showed some enrichment of sites of active translation co-localising with sites of viral RNA replication as determined by staining for dsRNA (Fig. 2E). However this enrichment fell short of the sequestration observed with vaccinia infection (David et al., 2012) with the majority of active translation localizing outside of the viral replication complexes.

**Figure 2.**
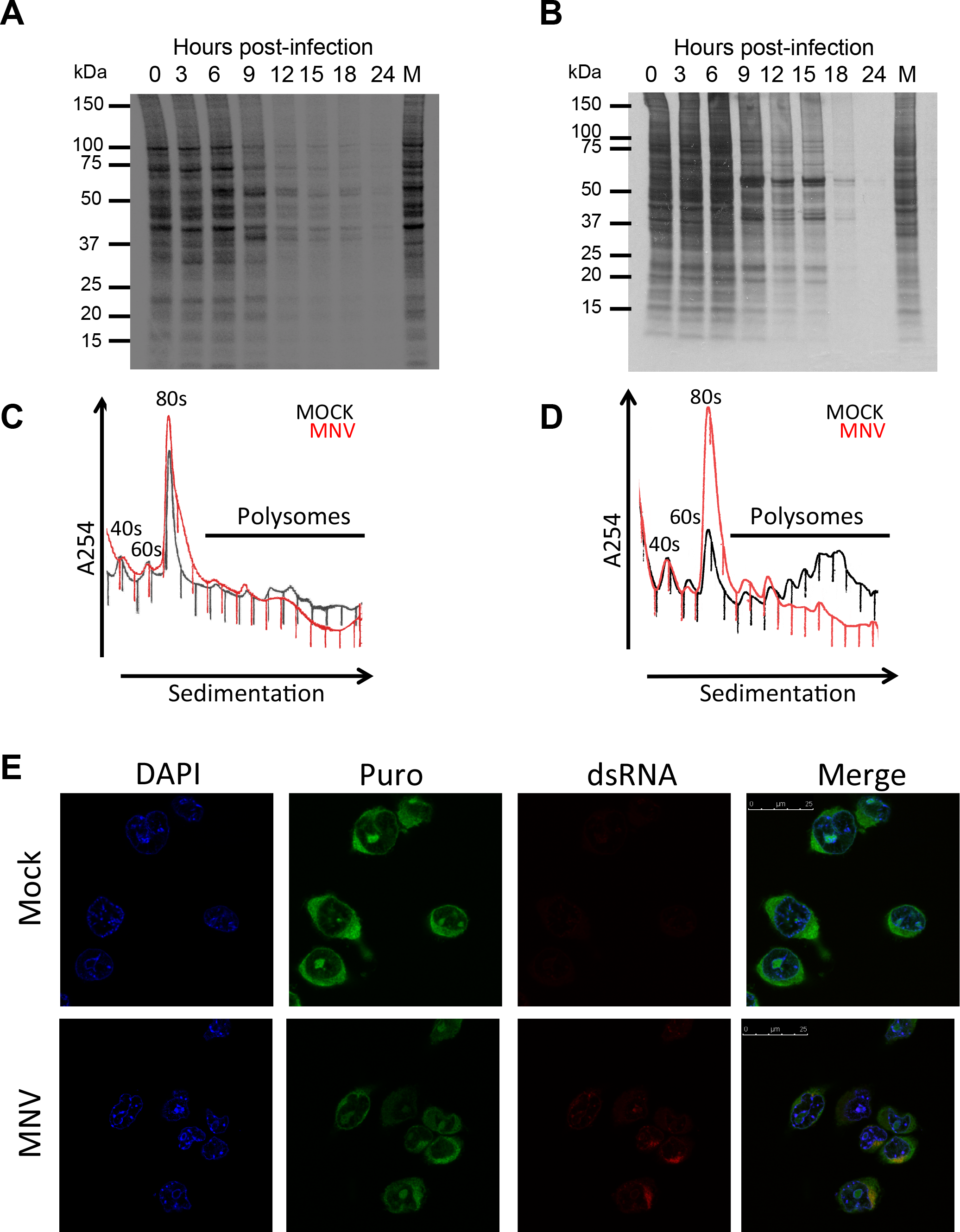
Norovirus infection alters the translational profile of the host cell. A) RAW 264.7 or B) BV-2 cells infected at an MOI of 10 TCID 50/cell with MNV were pulse labeled for 1h at the indicated timepoints and imaged on a phosphorimager. Polysome profiling of infected C) RAW 264.7 or D) BV-2 cells was performed. E) Puromycylation analysis of infected BV-2 cells shows some minor enrichment of sites of active translation co-localising with viral replication complexes visualized using anti-dsRNA.

### Quantitative proteomic analysis of MNV-infected cells and m7GTP-binding complexes reveals modifications to the eIF complex

We have previously described the novel mechanism of protein-primed VPg-dependent translation used by caliciviruses (Chaudhry et al., 2006a; Chung et al., 2014; Goodfellow et al., 2005; Hosmillo et al., 2014). Given the variation between the initiation factor requirements for host-cell mRNA and viral VPg-dependent RNA translation, a quantitative proteomics approach was used to investigate changes to both the level of relevant translation initiation factors in the host cell, as well as their ability to incorporate into the eIF complex at different times post infection. Using a stable isotope labeling approach (Munday et al., 2012; Ong and Mann, 2006; Ong et al., 2002), cells were labeled with light, medium or heavy stable isotope of arginine and lysine. Whole cell lysates were prepared from either mock or infected cells at 4 and 9 hours post infection and these samples used to determine the level of initiation factors within the cell. At the same time a m7GTP-sepharose enrichment was performed to determine the effect of viral infection on the composition of eIF4E-containing cap-binding complexes (Fig. 3A). Representative Coomassie staining (Fig. 3B) and western blot analysis (Fig. 3C) of the purified complex confirmed the enrichment of translation initiation factors, as well as recruitment of the viral VPg protein and loss of a marker for soluble cytoplasmic proteins (GAPDH).

**Figure 3.**
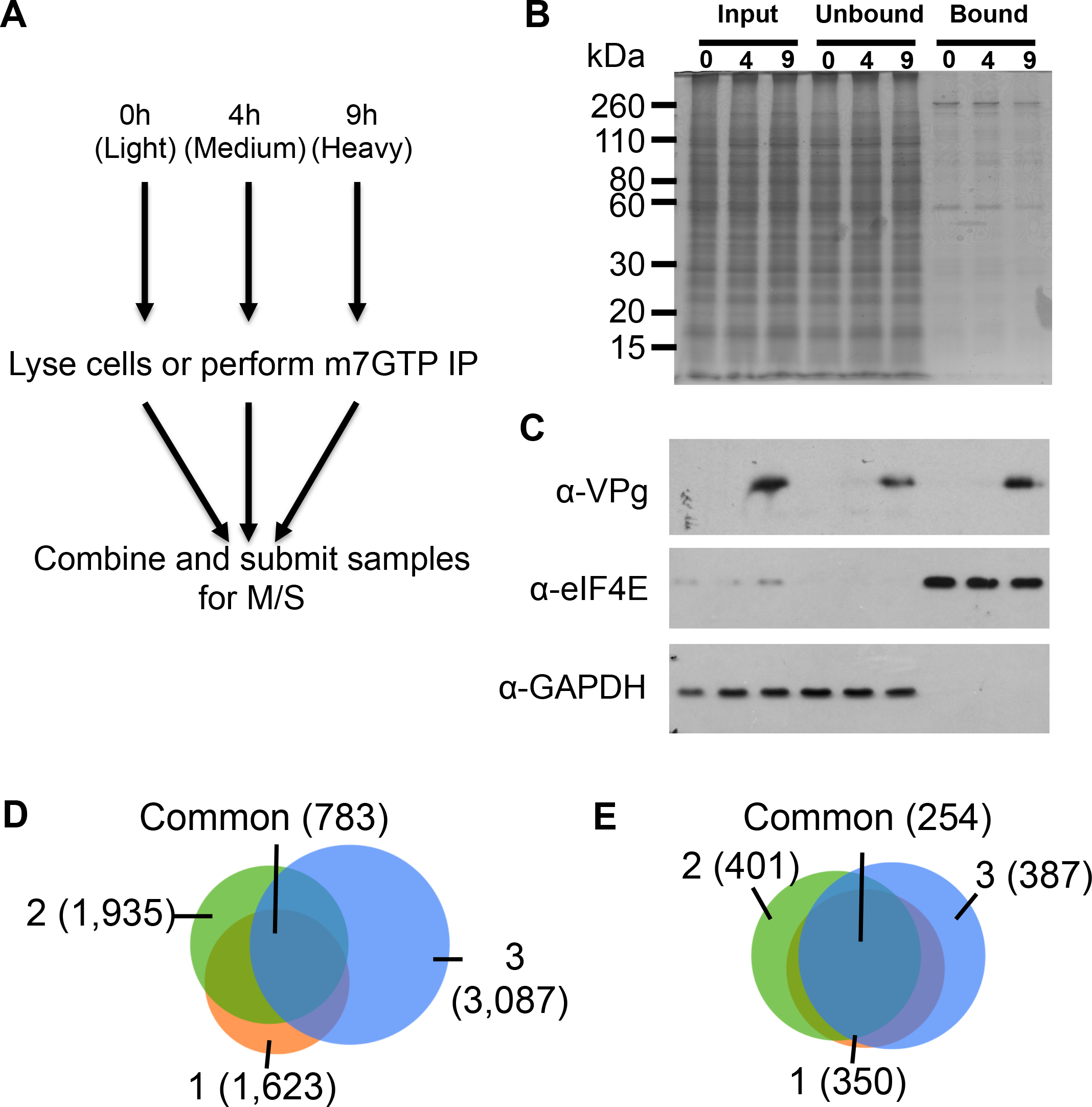
Quantitative proteomic analysis of translation initiation during MNV infection. SILAC-based quantitative proteomics was employed to investigate changes to eIF composition during high MOI MNV infection of BV-2 cells. The experimental layout is illustrated in A) with samples taken at mock (0h) early (4h) or late (9h) post-infection either lysed or subject to m7GTP-sepharose purification. A Coomassie gel (B) and representative western blots (C) confirm initiation factor enrichment. Venn diagrams illustrating experimental proteome coverage in D) whole cell lysate, or E) m7GTP-sepharose pulldowns.

Mass spectrometry analysis of independent biological replicates of the whole cell lysates identified 4203 proteins with two or more peptides that could be quantified (Fig 3D). As anticipated, the strategy used to enrich the eIF complex resulted in fewer proteins being identified across the three replicates (530), however a better overlap in the proteins quantified across the replicates (Fig 3E). Gene ontology analysis (Table S1) of the m7GTP-sepharose purified samples performed using STRING (Szklarczyk et al., 2015) revealed that proteins showing an arbitrary ≥2-fold change in their abundance at 9h post-infection were associated with translation initiation (p=2.85E-30) with changes to the eIF3 complex being particularly significant (p=3.12E-28). The complete proteomics dataset obtained from the analysis of both the whole cell lysate and the m7GTP-enriched complex is provided in tables S2 and S3.

Alterations to individual eIF components were assessed with no changes in the abundance of eIF4E within infected cells, or in its ability to bind to m7GTP-sepharose being observed (Fig. 4A). However changes to other components of the eIF4F complex were apparent. Reduced levels of the eIF4AII but not eIF4AI isoform of eIF4A were apparent at late time points in cells and in m7GTP-purified complexes (Fig. 4B). Another helicase, eIF4B, showed similar levels within whole cell lysates at late times post-infection, but was reduced within m7GTP-purified complexes (Fig. 4C). Isoform-specific variation in the impact of infection on eIF4G levels was also observed with eIF4GI showing slight, but not significant reductions in cellular abundance and m7GTP-sepharose binding, whereas eIF4GII showed both reduced overall abundance in cells and reduced association with eIF4E-containing complexes enriched on m7GTP-sepharose (Fig. 4D). As a component of the core of the eIF4F complex, eIF4G interacts with the small ribosomal subunit via the eIF3 complex, which was detected in its entirety in both the whole cell lysate and in the m7GTP-associated complex (Fig. 4E,F). The abundance of eIF3 components remained largely unaffected during infection (Fig. 4E), however at 9h post infection the ability of eIF4F to recruit the eIF3 complex was greatly diminished, consistent with the gene ontology analysis (Fig. 4F). Other proteins recruited to the eIF4F complex by eIF3 also showed a similarly reduced ability to bind m7GTP-sepharose and are detailed in figure S3A-D. Western blot analysis of eIF4E-containing complexes by m7GTP-sepharose enrichment from infected cells confirmed the loss of eIF3D and eIF4GII, as well as the impact of infection on eIF4AII expression (Fig. S3E).

**Figure 4.**
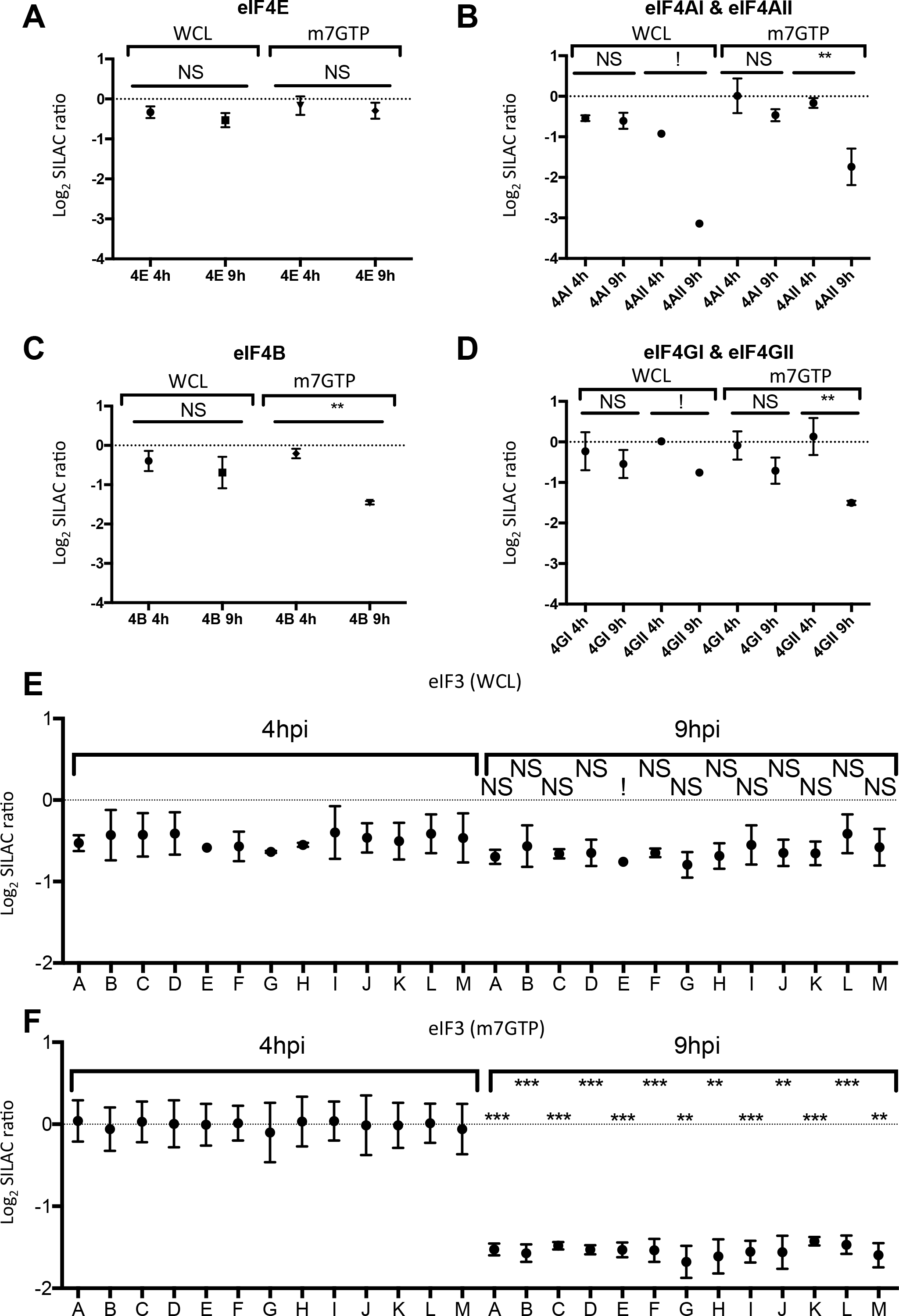
Norovirus infection alters the abundance and eIF4F association of cellular translation initiation factors. Mass spectrometry data for individual eIF components identified in the whole cell (WCL) or m7GTP-sepharose (m7GTP) experiments are shown, including A) eIF4E, B) eIF4A, C) eIF4B and D) eIF4G. The whole eIF3 complex was successfully identified by mass spectrometry and its relative abundance in E) WCL or F) m7GTP samples is shown. Significance was tested by t-test comparing changes to a control protein with unaltered abundance (eIF4E). (*=<0.05, **=<0.01, ***=<0.001, ****=<0.0001). Where insufficient replicates were identified to permit statistical analysis this is indicated with (!).

### The norovirus protease (NS6) contributes to translational inhibition via PABP cleavage

Previous studies with feline calicivirus (FCV) or recombinant norovirus protease suggested a potential role for the calicivirus 3C-like protease in the cleavage of PABP (Kuyumcu-Martinez M. et al., 2004). The biological consequence of the norovirus protease-mediated cleavage was not examined due to the lack of an available cell culture system at the time the study was undertaken. To determine if the MNV NS6 protease contributed to the loss of eIFs from the eIF4F complex, the ability of NS6 to cleave initiation factors was examined. Expression of a GFP-NS6 fusion protein resulted in the cleavage of PABP, whilst other initiation factors known to be targets of other 3C or 3C-like proteases remained uncleaved (Fig. 5A) (Belsham et al., 2000). Side-by-side comparison of the cleavage products of PABP from transfected cells with infected BV-2 or RAW 264.7 cells reveals that this cleavage product can be observed in infected cells, though additional cleavage products are also present (Fig. 5B). Analysis of time-course samples from infected cells shows the appearance of cleavage products from 9h post-infection, consistent with the timing of the impact on cellular translation (Fig. 5C). Not all of the PABP detected by western blot was cleaved during infection, in agreement with previous observations for other positive sense RNA viruses where cleavage of PABP has an impact on cellular translation (Kuyumcu-Martinez et al., 2002). The anti-PABP antibody used to examine cleavage during infection detects both PABP1 and PABP3, therefore it was not possible to distinguish if the partial cleavage observed by western blot was the result of an isoform specific effect of NS6. The expression of NS6 alone was sufficient to have a small but reproducible and significant impact on cellular translation in the absence of viral replication (Fig. 5D), without any obvious impact on cellular viability (Fig. S4B). Expression of NS6 alone in mock-or interferon-treated cells confirmed that NS6 expression alone was at least partially responsible for the reduced translation of induced ISGs (Fig. S4D,E). PABP consists of an N-terminal region containing multiple RNA recognition motifs and the eIF4G-binding site, and a C-terminal region containing the PABC-domain (Fig. S4A). Many viruses target the flexible linker region connecting these two domains (Smith and Gray, 2010). Mutational analysis was used to demonstrate that the NS6 protease cleavage site was Q440, also known as the 3Calt’ site (Fig. S4C). The biological consequence of PABP cleavage and the impact of cleavage on virus replication was examined by overexpressing either wild-type PABP or a non-cleavable (Q440A) form of PABP in MNV permissive cells. The expression of a non-cleavable form of PABP resulted in the partial restoration of ISG translation during infection (Fig. 5E) as low-levels of viperin induction was seen during infection. This partial restoration of ISG induction also resulted in a delayed replication during low multiplicity, multicycle replication typically causing a 1 log_10_ reduction in viral titres at 18h post-infection (Fig. 5F,G).

**Figure 5.**
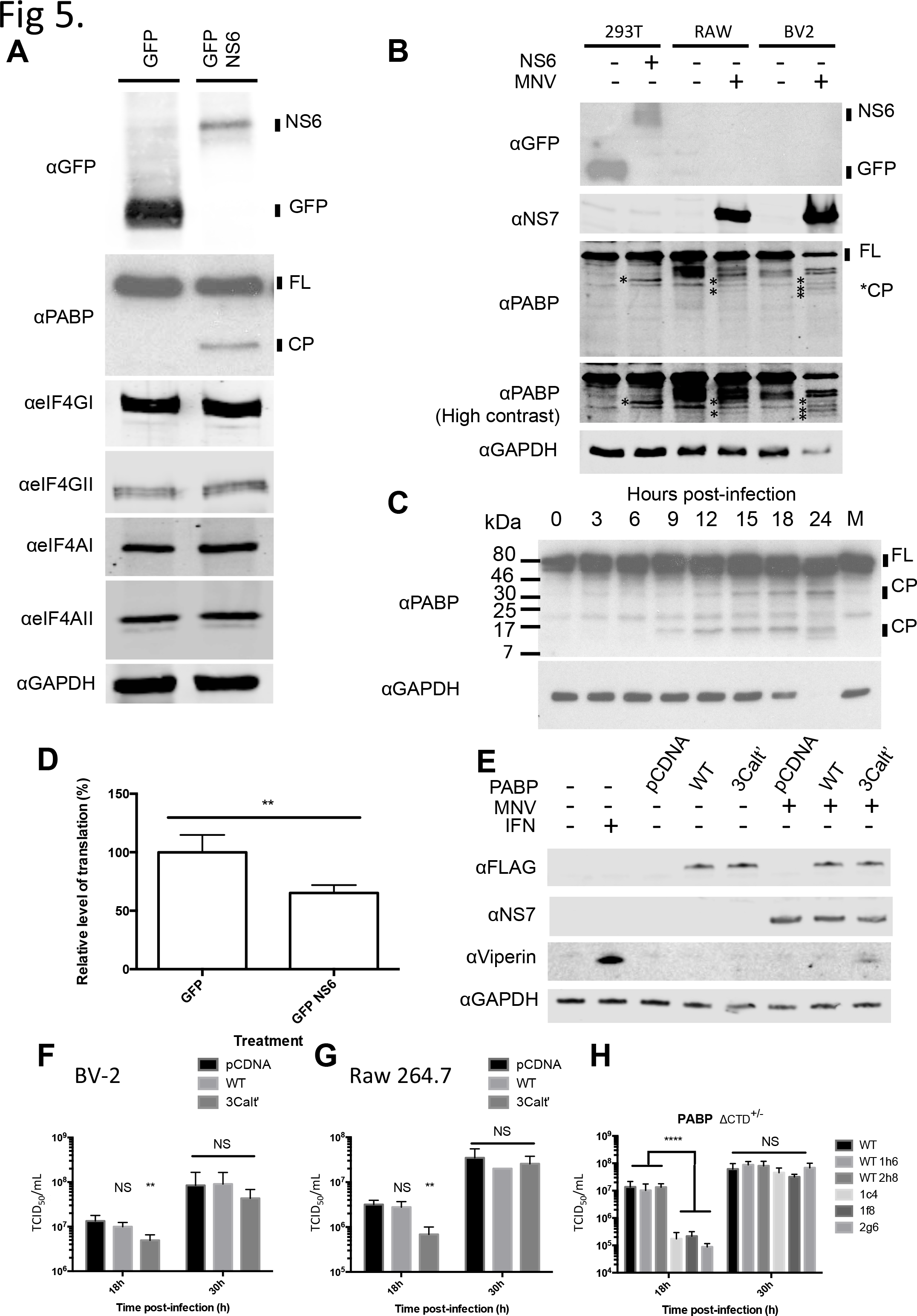
Cleavage of PABP by the norovirus protease NS6 contributes to reduced cellular translation. A) Western blot analysis of selected eIF proteins in cells transfected with the MNV protease NS6. B) Western blot analysis of NS6-cleaved PABP alongside lysates from infected RAW 264.7 and BV-2 cells C) Western blot analysis of PABP cleavage over a MOI 10 infection timecourse in BV-2 cells D) Analysis of global translation in 293T cells transfected with NS6 assessed by 35S-methionine pulse-labelling and quantification on a phosphorimager. E) Western blot analysis of cells transfected with wild-type or non-cleavable (Q440A) PABP. Viral titres obtained following low MOI (0.01) infection of F) BV-2 or G) RAW 264.7 cells transfected with wild-type or a non-cleavable form (Q440A) of PABP. H) Viral titres obtained following low MOI (0.01) infection of cells heterozygous for a truncated form of PABP. Error bars represent standard deviation. Statistical analysis was performed by one-way ANOVA. (*=<0.05, **=<0.01, ***=<0.001, ****=<0.0001).

To investigate whether infection or cleavage of PABP in infected cells impacted on PABP localization, confocal microsocopy was used to examine the localization of PABP in infected cells. Both mock and infected cells possessed diffuse cytoplasmic PABP localization, with no evidence of altered localization in response to MNV-infection apparent (Fig. S5A). Attempts were made to use CRISPR-mediated gene editing to generate MNV permissive cells expressing a non-cleavage form of PABPC1 (Fig S5B), however all clones that were isolated were heterozygous for a PABP C-terminal deletion (all clones possessed a frameshift at P433, resulting in a truncation 20 amino acids later) (Fig S5C). When not infected, cell viability (Fig. S5D) and translation (Fig. S5E) in these cells remained unaffected. Upon infection, an approximate 100-fold drop in viral titres at 18h post-infection was observed in cells containing the truncated form of PABP (Fig. 5H). These data demonstrate a role for NS6 cleavage of PABP in at least partially reducing cellular translation in infected cells.

To examine if NS6-mediated cleavage of PABP also resulted in the observed loss of eIF3 from the eIF4F complex the effect of NS6 cleavage on the recruitment of eIF3 to the eIF4F complex was examined by the enrichment of the eIF4F complex on m7GTP-sepharose. The recruitment of eIF3D to the eIF4F complex was not affected by PABP cleavage, yet both full length and the N-terminal cleavage products of PABP were recruited to the eIF4F complex (Fig. S4F). These data suggested that whilst NS6-mediated PABP cleavage plays a partial role in reducing cellular translation in MNV-infected cells, other pathways also contribute. Additional cleavage products of PABP in Fig. 5B may represent apoptotic cleavage products.

### Loss of eIF3 recruitment to translation initiation complexes is part of the cellular response to infection

Apoptosis is a key cellular pathway known to inhibit translation initiation (Clemens et al., 2000). MNV infection causes the induction of apoptosis through downregulation of survivin (Bok et al., 2009) leading to the activation of a number of caspases as well as cathepsin B (Furman et al., 2009a). The eIF4GI and II proteins are targets for caspase-mediated cleavage at multiple positions as shown in Fig. 6A (Clemens et al., 2000). The SILAC quantification presented in Fig 4 represents an averaged fold-change across all the quantified peptides from an individual protein. In the case of eIF4GI, over 300 peptides from this protein were identified in the m7GTP-sepharose purified samples across the entire protein, enabling the direct identification of eIF4G fragments that may remain associated with the eIF4F complex if caspase cleavage had occurred. Caspase cleavage of eIF4GI results in the production of three fragments, of which only the middle fragment (M-FAG) would be expected to bind efficiently to m7-GTP sepharose. Differing amounts of the N-FAG, M-FAG and C-FAG fragments were found associated with the eIF4F complex at 9 hours post infection, consistent with caspase-mediated cleavage resulting in the preferential loss of fragments that do not interact directly with eIF4E (Fig. 6B). Retention of N-FAG could be explained as an indirect interaction mediated through PABP and the mRNA To examine the correlation between induction of apoptosis and the impact of infection on cellular translation, the activation of caspase 3, the cleavage of PARP, eIF4GI and eIF4GII was examined by western blot. In BV-2 cells, loss of full-length eIF4GI and II and the appearance of prominent eIF4GII cleavage products were apparent from 9h post-infection, concomitant with the appearance of cleaved-caspase 3 and PARP (Fig. 6C). In contrast, during replication in RAW 264.7 cells the induction of apoptosis was less pronounced and somewhat delayed in agreement with previous observations (Bok et al., 2009; Furman et al., 2009b; McFadden et al., 2011) with only low levels of cleaved caspase being detected and incomplete cleavage of PARP (Fig. 6D). The marked cleavage of eIF4GI and II was also not readily observed. Cleaved caspase 3 and PARP cleavage were observed from 9h post-infection, however unlike in BV-2 cells, levels of cleaved caspase 3 remained low, and PARP cleavage did not reach completion. Whilst the contribution of individual caspases can vary, efficient PARP cleavage is a hallmark of apoptotic cell death. Incomplete PARP cleavage has been previously linked to accelerated cell death through a combination of apoptosis and necroptosis and linked to low intracellular ATP and NAD+ levels (Herceg and Wang, 1999). This suggests that cell death in RAW264.7 cells may not be exclusively apoptotic and offers an explanation as to why the apoptotic phenotype is less pronounced in these cells.

**Figure 6.**
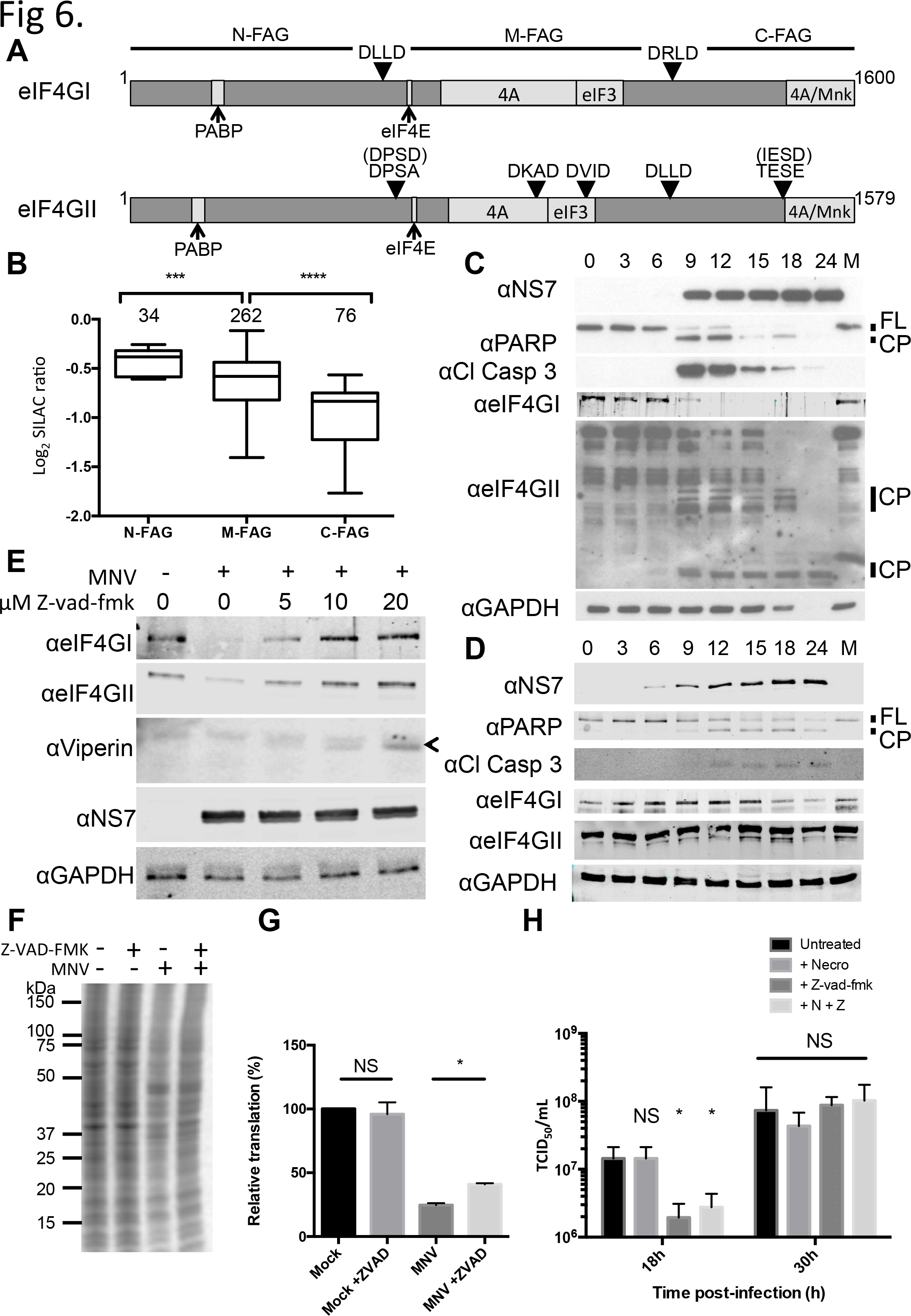
Induction of apoptosis and caspase cleavage of eIF4F components also contributes to altered translation and alters eIF4F composition. A) A diagram illustrating the structure of eIF4GI and II as well as their caspase cleavage sites. B) Quantification of peptides mapping to the N-FAG, M-FAG or C-FAG domains of eIF4GI binding m7GTP-sepharose beads at 9h post-infection. C) Western blotting against eIF4G and markers of apoptosis for an infection timecourse from BV-2 or D) RAW 264.7 cells. E) Western blotting of eIF4GI or II, and Viperin in the presence of varying amounts of the caspase inhibitor z-vad-fmk. The specific viperin band is highlighted with ‘<’ F) ^35^S-Methionine pulse labeling of z-vad-fmk-treated BV-2 cells Mock-or Infected with MNV at 9h postinfection. G). Quantification of ^35^S-methionine labeled cells showing restoration of translation following z-vad-fmk treatment H). Viral titres obtained following low MOI (0.01) infection of BV-2 cells treated with z-vad-fmk (20µM), necrostatin-1 (40µM) singly or in combination. Error bars represent standard deviation. Statistical analysis was performed by one-way ANOVA. (*=<0.05, **=<0.01, ***=<0.001, ****=<0.0001).

To determine if the cleavage of eIF4GI and II during MNV infection was the result of apoptosis, the effect of the pan-caspase inhibitor z-vad-fmk on eIF4G cleavage was examined. As previous studies have indicated that the inhibition of apoptosis in BV-2 cells results in rapid induction of necroptosis (Fricker et al., 2013), the necroptosis inhibitor necrostatin-1 was included where noted. eIF4GI and II levels were restored by the inhibition of caspases in a dose-dependent manner which also correlated with a partial restoration of translation of the representative ISGs (Fig. 6E). The impact of infection on the cellular translation profile of cells was also partially restored when apoptosis was inhibited (Fig. 6F). Furthermore, the inhibition of apoptosis and the resulting increase in the translation of induced ISG mRNAs lead to a delayed replication during multicycle replication (Fig. 6H).

## Discussion

As obligate intracellular pathogens, viruses must replicate within host cells and must therefore balance the use of host resources and the ability to evade/control the cellular response to infection. In this study we demonstrate that noroviruses alter cellular translation through the modification of translation initiation factors to not only favour viral translation but also impede the translation of genes induced as a result of the innate immune response. This offers a new mechanism by which norovirus is able to regulate the immune response, and demonstrate that by modifying translation in infected cells, noroviruses are capable of preventing production of ISGs such as STAT1 and ISG15 known to be important for control of norovirus infection (Karst et al., 2003; Rodriguez et al., 2014).

In the current study we observed that PABP cleavage by the norovirus protease plays a role in modification of host translation and can impact on the translation of induced ISGs. PABP cleavage has also been observed following infection with a range of viruses including other caliciviruses (Kuyumcu-Martinez M. et al., 2004), picornaviruses, HIV (Alvarez et al., 2006; Castelló et al., 2009a) and others (Smith and Gray, 2010). The actual mechanism by which PABP cleavage inhibits or modifies cellular translation remains unclear. PABP plays multiple roles within the cell; cytoplasmic PABP is involved in translation enhancement via a bridging interaction with eIF4G leading to a “closed loop” conformation of the RNA which in turns is believed to increase RNA stability and promote ribosome recycling (Wells et al., 1998). Through additional interactions with the ribosomal release factor eRF3, PABP is also believed to play a role in translation termination contributing to the control of nonsense-mediated mRNA decay (Amrani et al., 2004; Uchida et al., 2002). Whilst early research hypothesized that cleavage would prevent mRNA circularization (Joachims et al., 1999), more recent data suggests that the N-terminal region of PABP is necessary and sufficient for both poly(A) and eIF4G-binding (Kuyumcu-Martinez N. M. et al., 2004). The cleavage of PABP by picornaviruses is also incomplete and targets polysome associated PABP (Kuyumcu-Martinez M. et al., 2004; Kuyumcu-Martinez N. M. et al., 2004; Rivera and Lloyd, 2008) indicating the protease targets only a subset of PABP molecules. The C-terminus of PABP is required for binding of a number of protein partners including PAIP1 and 2, and eRF3 (Hoshino et al., 1999; Kozlov et al., 2001). It is likely that cleavage inhibits the recruitment of these proteins to actively translating mRNA, leading to reduced ribosome release and recycling of subunits (Kuyumcu-Martinez N. M. et al., 2004). The norovirus NS6-mediated cleavage removes the C-terminal oligomerization domain, which could act in a dominant manner to prevent the extension of PABP oligomers on the polyA tail of cellular (or viral) mRNA (Kühn and Pieler, 1996). However the impact of limited PABP cleavage on the oligomerization of PABP on the poly A tail is yet to be fully explored. Of note, our data from heterozygous CRISPR-modified cells show that a mixed population of full length PABP and the C-terminally truncated PABP fragment are sufficient for growth and normal levels of translation in unstressed/uninfected cells. This suggests that partial recruitment of PABP-binding proteins to polysomes is sufficient for normal protein synthesis, however more complete loss is inhibitory (Fig. 7B).

**Figure 7.**
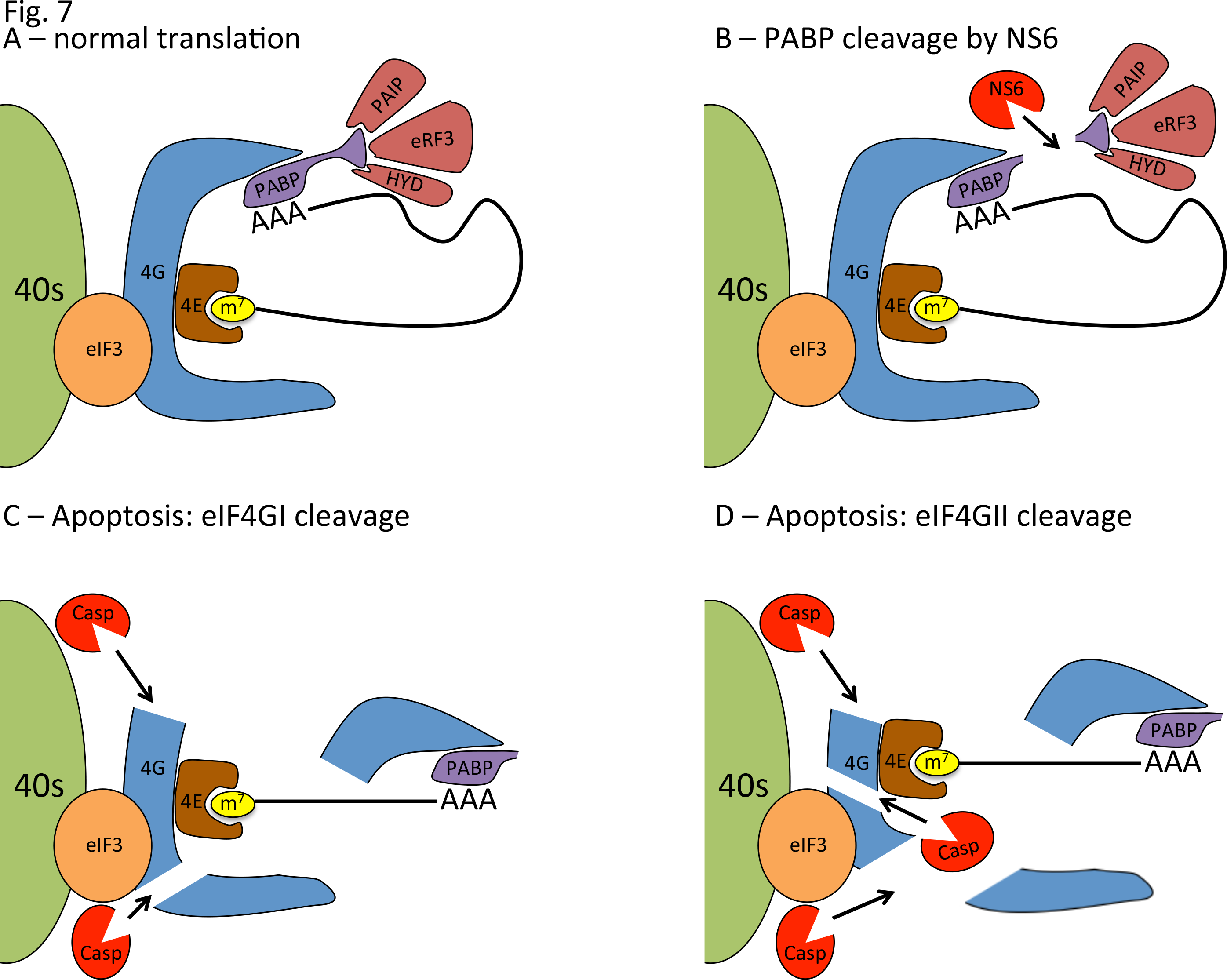
Model for modifications to eIF4F during norovirus infection. A)In healthy cells, the 5’ cap of mRNA is bound by eIF4E. This is bound by the scaffolding protein eIF4G which allows binding to other initiation factors (eIF4A, eIF3, PABP) as well as recruitment to the ribosome via the eIF3 complex. The norovirus protease NS6 alone can inhibit translation through cleavage of PABP, though larger scale alterations to the initiation factor complex are not observed. In infected cells, the induction of apoptosis can result in further modification of the eIF complex with caspase cleavage of C) eIF4GI separating the eIF4E and PABP-binding regions and abolishing the circularization of translating mRNAs. D) Cleavage of eIF4GII by caspases is more extensive and in addition to the effects observed with eIF4GI cleavage, separates the eIF4E and eIF3-binding domains of eIF4GII, preventing recruitment of mRNA to the ribosome.

Caspase-mediated cleavage of initiation factors has multiple effects on cellular translation (Clemens et al., 1998; Marissen and Lloyd, 1998)(Marissen et al., 2000). In the case of eIF4GI, cleavage causes the linearization of the mRNA as the PABP-binding N-FAG fragment of eIF4GI and the middle M-FAG fragment containing the eIF4E and eIF3-binding sites are separated following cleavage (Fig. 7C). Apoptotic translation shutoff correlates with cleavage of eIF4GII where additional caspase target sites further degrade eIF4GII and separate the eIF4E and eIF3 binding sites preventing recruitment of the mRNA to 43s subunits (Fig. 7D) (Marissen et al., 2000). Of note, VPg-dependent viral translation appears to continue even after apoptotic cleavage of eIF4G has initiated (Fig. 2,6). This is consistent with recent data on VPg-dependent translation where the middle fragment of eIF4G is sufficient for norovirus translation (Chung et al., 2014; Leen et al., 2016). The importance of this late viral translation for virus biology remains to be determined.

The mass spectrometry results successfully identified modification consistent with caspase-cleavage of eIF4G, but also yielded further details on alterations to the eIF complex, including diminished eIF4AII, but not eIF4AI levels, and alterations to eIF4B binding. Particularly in the case of eIF4AII, there have been conflicting reports that this specific form of eIF4A is vital for microRNA-mediated translation inhibition (Galicia-Vázquez et al., 2015; Meijer et al., 2013). Whether the loss of eIF4AII is due to the viral or host response merits further investigation. This work represents the first use of SILAC-based quantitative proteomics to study norovirus infection or virus-mediated translation inhibition. Furthermore, this study makes clear that quantitative proteomics is capable of offering up a new level of detail with regards to identifying the roles and responses of individual eIF components under stress conditions, and their contribution to the translation efficiency of individual mRNAs.

Other viruses have been shown to modify eIF components in order to alter cellular translation (Mohr and Sonenberg, 2012). One example would be African Swine Fever Virus which utilizes both eIF4E phosphorylation and redistribution of the translational machinery to viral factories in order to control host translation and the immune response (Castelló et al., 2009b). However, the best characterized example of eIF4F modification is from the picornaviruses which utilize two viral proteases to create a environment where host translation is completely inhibited (Gradi et al., 1998). The effects observed in norovirus-infected cells in many ways resemble those generated in picornavirus infection, however there are important distinctions. Firstly, the scale of inhibition seen is clearly different with picornaviruses causing complete host shutoff, whilst noroviruses merely cause a reduction of host translation (Fig. 2). Secondly, shutoff in picornavirus infection is an early event and allows the exclusive use of host translation apparatus for efficient viral translation (Gradi et al., 1998), whereas in norovirus infection, altered translation is a relatively late event and comes at a time when viral replication appears largely complete (Fig. 1,2). Whilst the shutoff observed in picornaviruses would also inhibit production of ISGs in response to infection, in norovirus infection this would appear to be the primary effect of reduced cellular translation at this time during infection (Fig. 1,2). A final distinction is in the mechanism of inhibition, with picornavirus shutoff driven entirely by two viral proteases (2A, 3C) that are necessary and sufficient for the observed phenotype. In contrast norovirus utilizes just a single viral protease-NS6, a 3C-like protease which of the initiation factors tested, cleaves only PABP (Fig. 5). However, norovirus infection is capable of inducing further alterations to translation initiation by utilizing the induction of apoptosis and in particular, caspase activation in order to mimic the effects of 2A protease cleavage on eIF4G (Fig. 6)(Lamphear et al., 1995). Whilst apoptosis is also induced in poliovirus-infected cells, the contribution of this to eIF4G cleavage is unclear (Calandria et al., 2004). The cells used in this study are macrophage or microglia-like cell lines, and in the infected host the virus is known to be able to infect dendritic cells and macrophages (Wobus et al., 2004), and most recently, B cells (Jones et al., 2014). Whilst the pro-viral nature of the apoptotic response in norovirus infection has been discussed previously, it is possible that this mechanism plays a role in a subset of infected cells within the host. Of particular note is the recent finding that enteric bacteria may also be able to modify this response, with Salmonella co-infection inhibiting MNV-induced apoptosis, diminishing viral replication and notably also increasing cytokine levels produced in response to infection (Agnihothram et al., 2015), fitting with our hypothesis that apoptosis is used by noroviruses as a mechanism to suppress the translation of induced ISGs.

In summary this study demonstrates than norovirus infection modifies the ability of host cells to respond to infection by limiting the translation of induced mRNAs. The alterations to host translation observed are induced both directly by the virus, as well as through the induction of apoptosis, and serve to counter the paracrine impact of the innate response.

## Experimental Procedures

### Cells and Viruses

Murine RAW 264.7 and BV-2 cells were used for infection experiments; human HEK-293T cells were used for the indicated transfection experiments. All infections were performed with murine norovirus strain CW1 (Chaudhry et al., 2007). All titre calculations were performed by TCID_50_. All infections were performed at high MOI (10 TCID_50_/cell) unless explicitly indicated, in which case a low MOI of 0.01 TCID_50_/cell was used. See Supplemental Experimental Procedures for more details.

### Cell lysis and ^35^S Methionine labeling

Cells were lysed in RIPA buffer for analysis of whole cell extracts. For metabolic labeling experiments utilizing ^35^S-Methionine media was replaced with DMEM containing ^35^S-methionine 30 minutes prior to the indicated time and the samples harvested 30 minutes after the indicated time in RIPA buffer.

### m7GTP-sepharose enrichment and polysome profiling

For analysis of the eIF complex, initiation factors were enriched on m7GTP-sepharose beads (Jena Biosciences) following lysis in m7GTP lysis buffer as described in Chung et al., 2014. Polysome profiling was accomplished by centrifuging cytoplasmic lysates for 90 min at 200,000 × g over a 10-50% sucrose gradient and analysed using an Isco Fractionator measuring absorbance at 254nm. For full details see Supplemental Experimental Procedures.

### Mass spectrometry analysis

Cells were grown in DMEM containing stable isotope labeled forms of arginine and lysine for 5 passages, with labeling confirmed by mass spectrometry. Samples were harvested and combined at the indicated timepoints following either lysis or m7GTP-sepharose enrichment. These samples were subject to SDS-PAGE electrophoresis, and processed by in-gel trypsinisation followed by LC-MS/MS analysis on a Orbitrap Velos instrument at the University of Bristol. Results were analysed using Thermo Proteome Discoverer software. Full details are given in Supplemental Experimental Procedures.

### qRT-PCR

RNA samples were extracted using the Genelute mammalian total RNA extraction kit (Promega) and qRT-PCR analysis performed by the SYBR green method on a Viia 7 instrument. Relative quantification was performed by the ΔΔCt method relative to a GAPDH standard, and absolute quantification was performed by comparing RNA copies to a serially diluted DNA standard.

### Western blotting analysis

Cell lysates were subject to SDS-PAGE electrophoresis and transferred to nitrocellulose membranes according to standard protocols. Blocking and antibody incubation steps were performed in 5% BSA or 5% non-fat dried milk as appropriate for the antibody. Detection was performed using either HRP-conjugated antibodies and chemiluminescent detection, or infrared-dye-conjugated secondary antibodies and detection on a Li-Cor Odyssey imager. All densitometry was performed on samples analysed on a Li-Cor Odyssey imager using the ImageStudioLite software (Li-Cor). Full experimental and antibody details are given in Supplemental Experimental Procedures.

### Immunofluorescence microscopy & puromycylation

Puromycylation was performed as described in (David et al., 2012). In brief, cells grown on glass coverslips were incubated for 5 minutes in cell culture media supplemented with puromycin and emetine at 37°C. Cells were transferred to, and maintained on ice for subsequent extraction steps. Cells were incubated for 2 min with permeabilization buffer. Cells were then washed once with polysome buffer, and fixed with PFA for 15 min at room temperature. PFA was aspirated, PBS was added, and cells were maintained at 4°C. Subsequent antibody incubations and washing steps followed standard protocols and are described in (Vashist et al., 2012). Imaging was performed on a Leica Sp5 confocal microscopy using a 63× oil objective. Image analysis was performed in the Leica Lita software (Leica Microsystems). Full details are given in Supplemental Experimental Procedures.

### Accession numbers

The mass spectrometry proteomics data have been deposited to the ProteomeXchange Consortium via the PRIDE (Vizcaíno et al., 2016) partner repository with the dataset identifiers PXD004015 (whole cell SILAC experiments) and PXD004019 (m7GTP enrichment).

## Supplemental Information

Supplemental information associated with this manuscript includes Supplemental Experimental Procedures, Five Figures, and Three Tables.

## Author contributions

EE, FS, SC, SV, SS, RL, KH and IG conceived and designed experiments. EE, SC, FS, SV, SS, KH performed experiments. SV and RL provided reagents. EE and IG wrote the manuscript. All authors gave final approval of the manuscript.

## Acknowledgements

This work was supported by grants from the Wellcome Trust (097997/Z/11/Z) and BBSRC (Refs: BB/N001176/1 and BB/K002465/1) to IG. RL is support bi a grant from the National Institutes for Health of the United States of America (AI50237). IG is a Wellcome Senior Fellow. Dr Ian Brierley (University of Cambridge) and Dr Mark Rodgers (Cambridge Bioscience) are thanked for their assistance with polysome profiling. This work was also supported by the Cambridge NIHR BRC cell phenotyping hub, and in particular their assistance with microscopy.

## Supplemental Figure Legends

**Figure S1.**
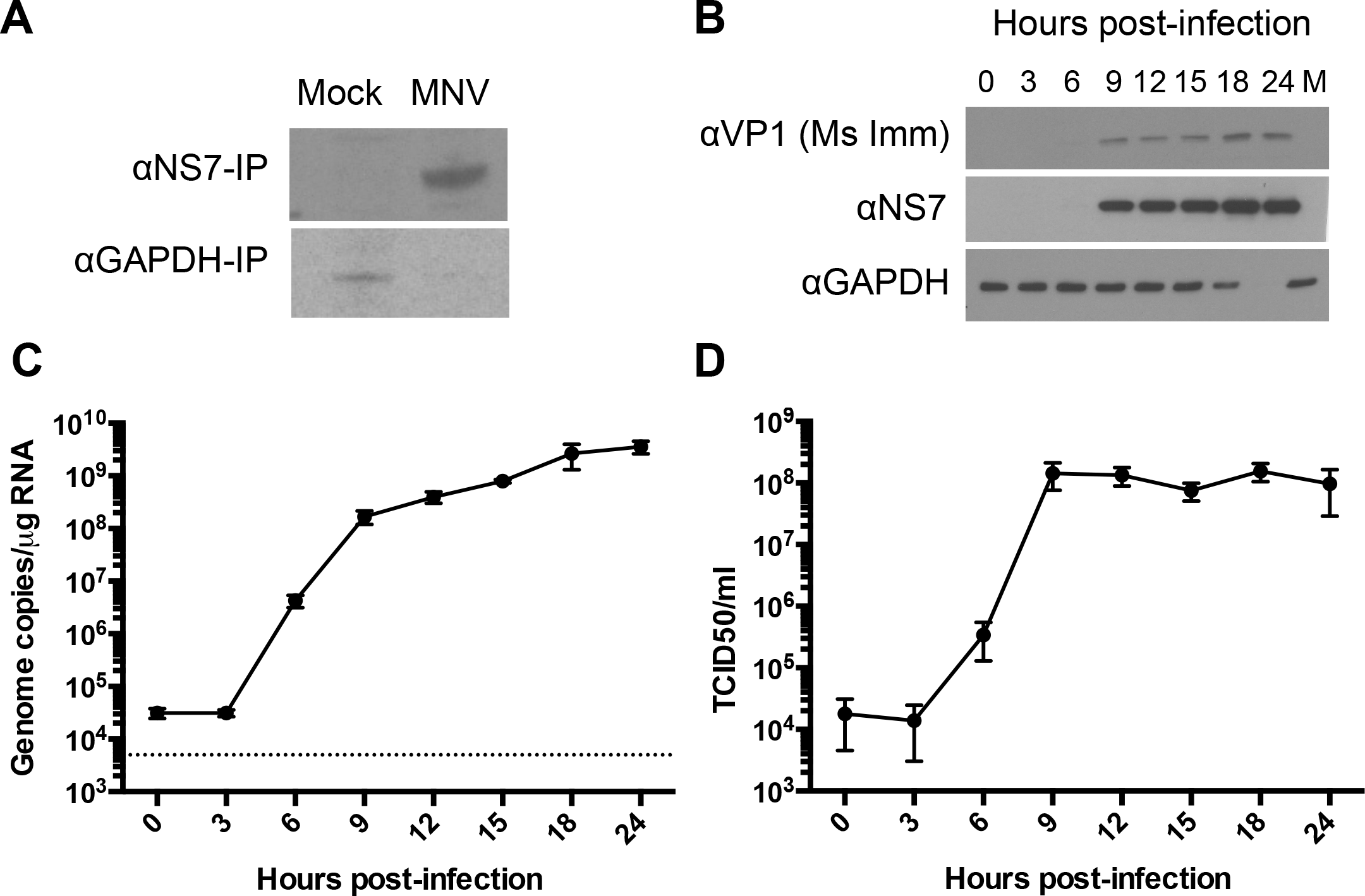
Translation at late times post-infection is dominated by viral proteins. A) Immunoprecipitation of viral (NS7) and cellular (GAPDH) proteins from S35-labelled infected BV-2 cells at 12hpi reveals translation of viral, but not cellular proteins. Analysis of a high multiplicity of infection timecourse in BV-2 cells shows viral B) protein, C) genome copies and D) titres have largely peaked by 9 hpi.

**Figure S2.**
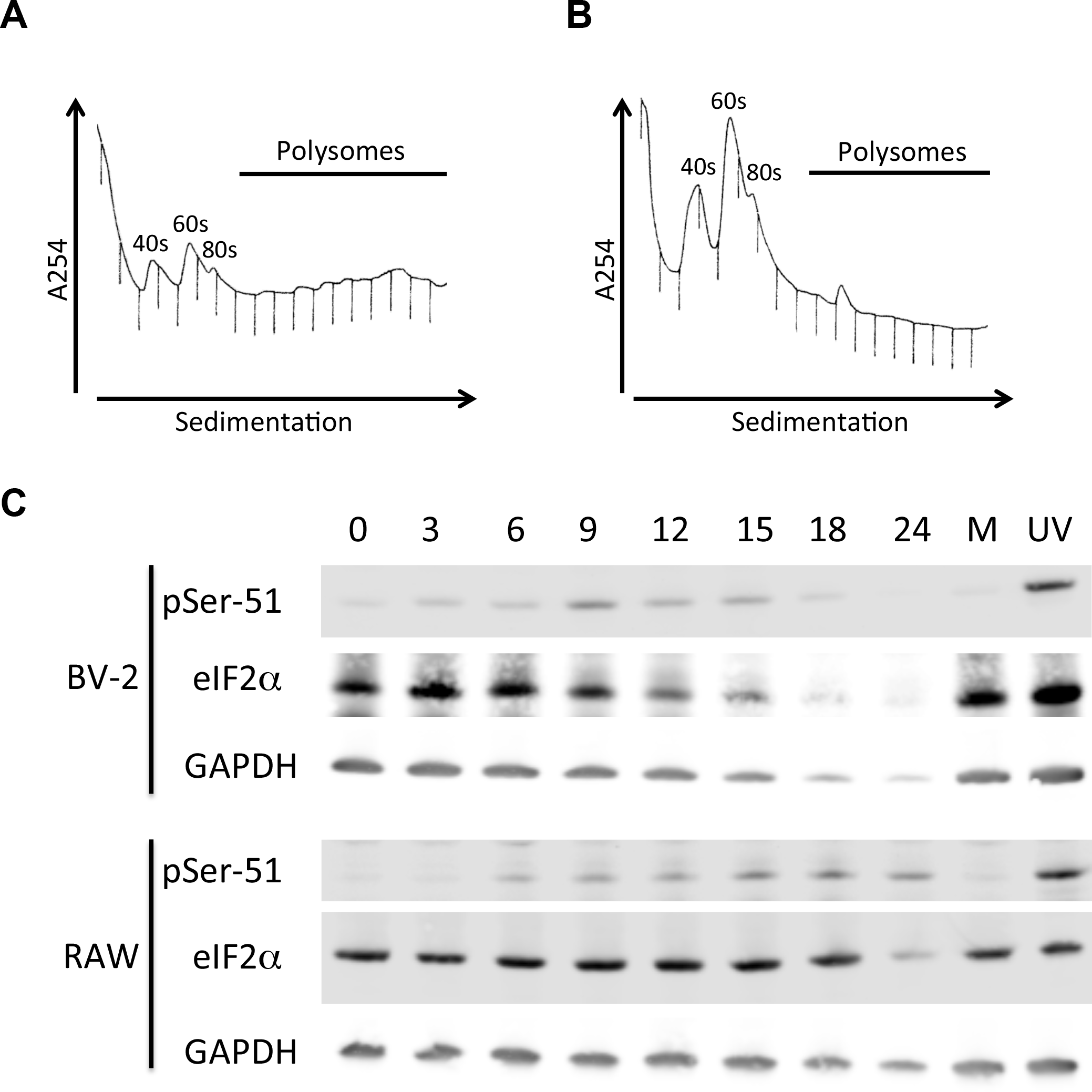
A defect in translation inhibiton occurs without high eIF2α phosphorylation or sequestration of eIF components. Polysome profiling of A) Mock or B) MNV-infected BV-2 cells performed under high salt conditions (400mM KCl). C) Western blot analysis of eIF2α phosphorylation in MNV-infected RAW 264.7 or BV-2 cells **(Associated with Figure 2.)**.

**Figure S3.**
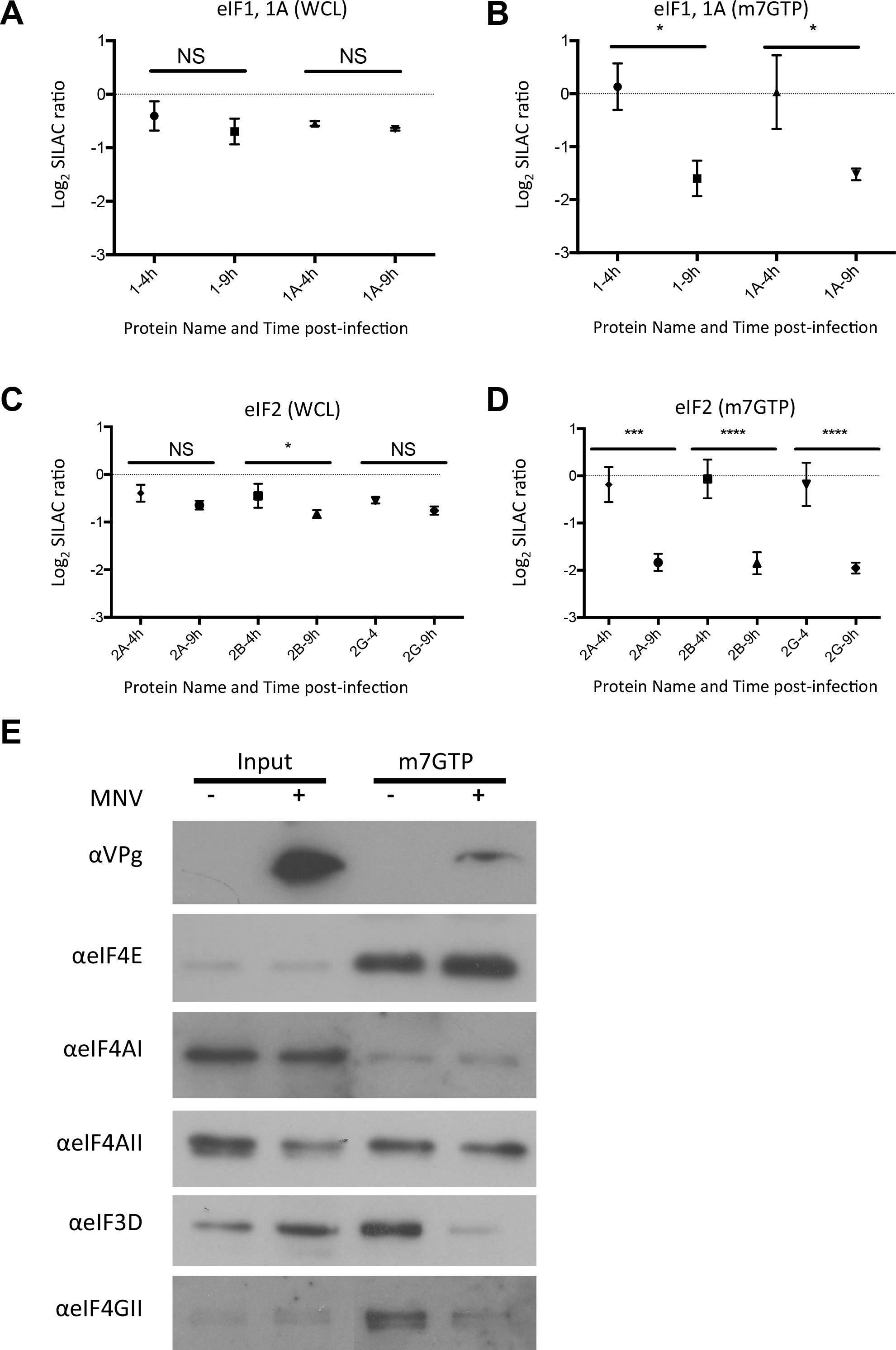
eIF changes upon MNV-infection and validation by western blotting. Downstream of eIF3, additional initiation factor components also show diminished binding to m7GTP-sepharose at late times following MNV-infection, including A-B) eIF1, and C-D) eIF2. E) Western blotting analysis of selected initiation factor binding to m7GTP-sepharose identified in mass spectrometry analysis. Significance was tested by t-test comparing changes to a control protein with unaltered abundance (eIF4E). (*=<0.05, **=<0.01, ***=<0.001, ****=<0.0001). Where insufficient replicates were identified to permit statistical analysis this is indicated with (!).**(Associated with Figure 4.)**

**Figure S4.**
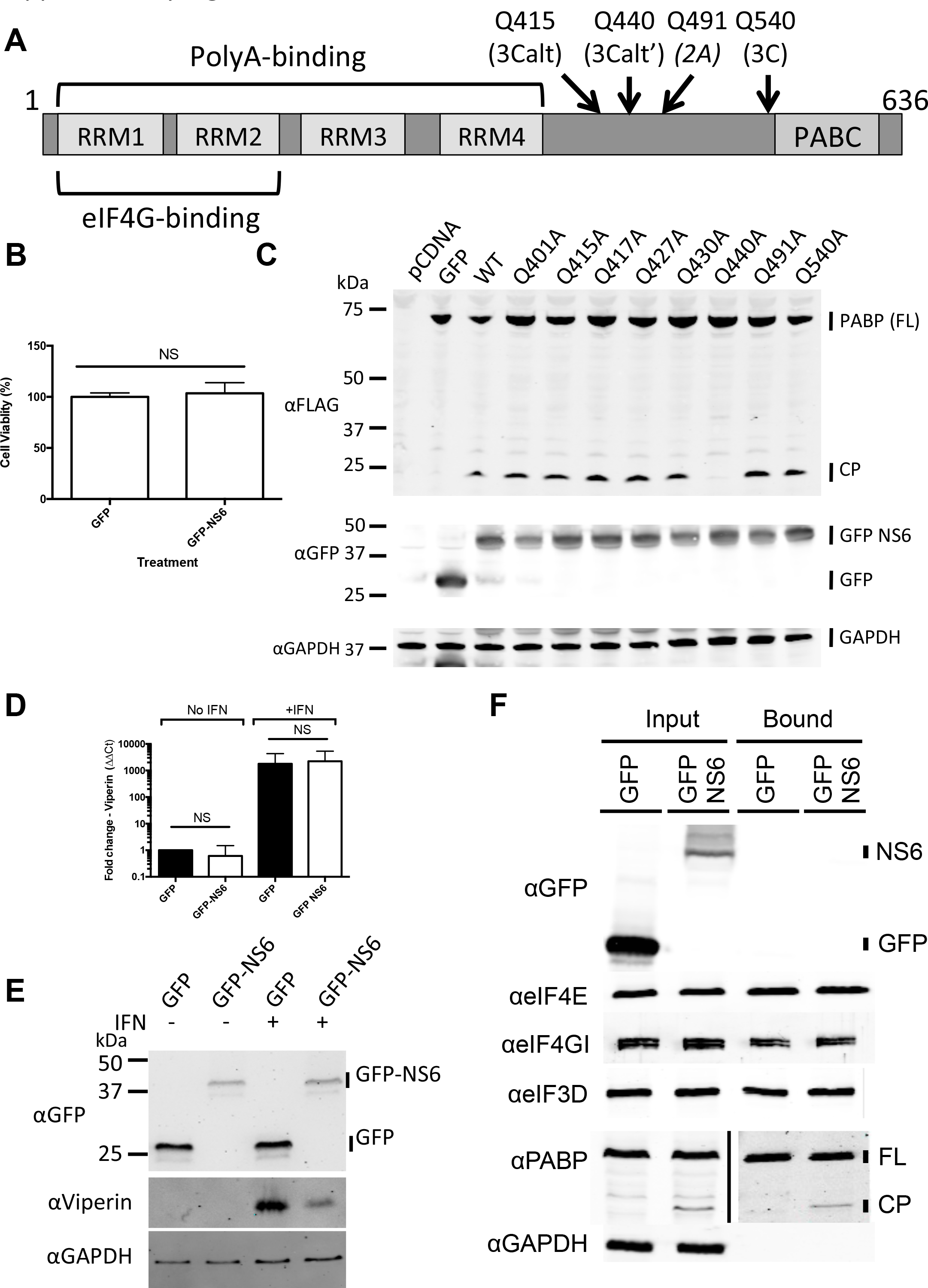
Identification of PABP cleavage sites and contribution to eIF complex modification and phenotype. A) A schematic illustration of the domains within PABPC1 protein (Uniprot accession P29341). B) Cell viability assay using Cell Titre Blue in cells transfected with pEGFP-C1 NS6. C) Western blot analysis of FLAG-tagged WT or mutant PABP D) qRT-PCR or E) western blot analysis of the levels of the ISG Viperin in pEGFP-C1 NS6 transfected 293T cells F) Western blot analysis of m7GTP-sepharose pulldowns from 293T cells transfected with pEGFP-C1 NS6. Error bars represent standard deviation. **(Associated with Figure 5.)**

**Figure S5.**
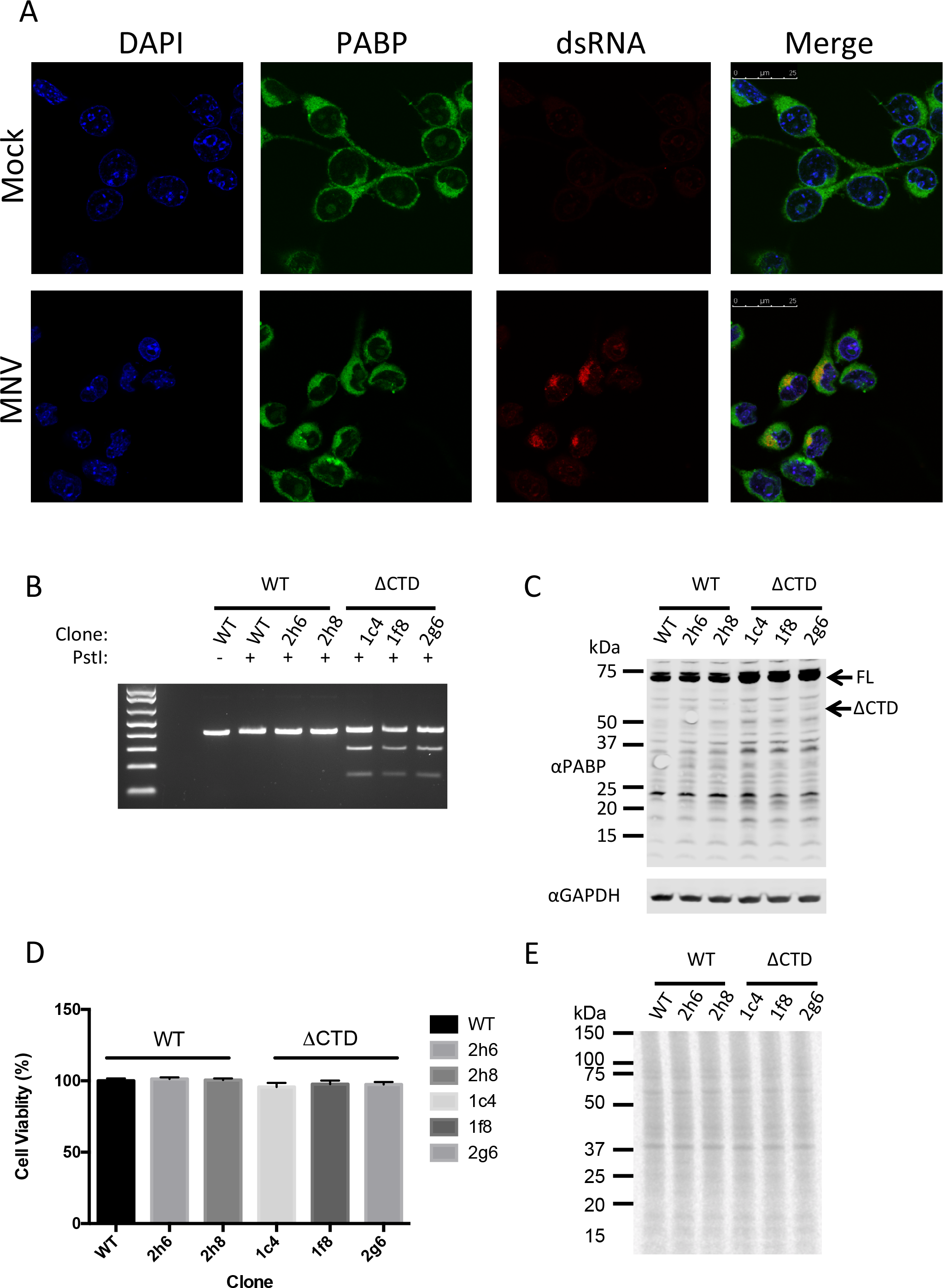
Confocal microscopy of PABP localization & generation of PABP ΔCTD (+/−) BV-2. A) Confocal microscopy of BV-2 cells at 12h post-infection with MNV showing no alteration in PABP localization. B) Amplicons from cells heterozygous for a PABP ΔCTD mutation contained a *PstI* restriction site introduced by homologous recombination. C) Western blot analysis of wild-type or PABP ΔCTD. D) Cell titre blue cell viability assay and E) translation from uninfected wild-type or PABP ΔCTD heterozygous BV-2 cells was comparable. **(Associated with Figure 5.)**

## Table legends

**Table S1. STRING analysis of proteins showing a ≥2-fold change in abundance in the 9h m7G-sepharose pulldown.** Proteins showing a >2-fold alteration in m7GTP-sepharose binding were inputted into the STRINGv10 database (Szklarczyk et al., 2015). Enriched GO Processes, Molecular Functions, and Cellular compartments are shown. **(Associated with Figure 3.)**

**Table S2. Whole cell lysate SILAC data from MNV-infected BV-2 cells.** Combined SILAC data from the three experimental repeats. The uniprot accession number, number of unique peptides from each experiment, and log_2_ ratios for each condition are listed. **(Associated with Figure 3.)**

**Table S3. m7GTP-sepharose pulldown SILAC data from MNV-infected BV-2 cells.** Combined SILAC data from the three experimental repeats. The uniprot accession number, number of unique peptides from each experiment, and log2 ratios for each condition are listed. **(Associated with Figure 3.)**

